# Virus Induced Gene Silencing in *Calendula officinalis* (pot marigold)

**DOI:** 10.64898/2026.02.16.706131

**Authors:** Daria Cuthbert, Connor Tansley, Melissa Salmon, Nicola Patron

## Abstract

Virus induced gene silencing (VIGS) is a method that exploits plant antiviral defence mechanisms to downregulate endogenous genes. The technique is versatile, rapid, and widely used for functional genomics studies. Here we report a method for VIGS in the medicinal plant, *Calendula officinalis* (pot marigold). This species produces anti-inflammatory triterpenoids and has also been bred and cultivated as an ornamental plant. We describe a method for the injection of *Agrobacterium tumefaciens* cultures into leaf midribs and compare visual marker genes for tracking VIGS utilising constructs that simultaneously target visual marker and target genes. We use these tools to demonstrate that silencing a gene encoding cycloartenol synthase results in changes to leaf phytosterols. This method could be used to further investigate the genetic basis of specialised metabolism in this species and could be adapted to other members of the Asteraceae family, many of which are of economical and chemical value.

## Introduction

RNA silencing is a key mechanism for regulating gene expression in eukaryotes in which epigenetic modifications are made to DNA sequences (transcriptional gene silencing; TGS) or messenger RNAs (mRNAs) are degraded or prevented from translating (post transcriptional gene silencing; PTGS) (Zhang et al., 2015; Zhan and Meyers, 2023). In plants, RNA silencing is also used for antiviral immunity. In this process, viral double-stranded RNAs (dsRNAs) are cleaved into small interfering RNAs (siRNAs) that bind to members of the argonaute (AGO) protein family and are incorporated into RNA-induced silencing complexes (RISC) where they serve as guides for the cleavage or translational suppression of viral RNAs (Lopez-Gomollon and Baulcombe, 2022). This process, known as virus-induced gene silencing (VIGS), has been adapted to enable the silencing of endogenous genes for functional studies and biotechnological applications.

VIGS typically employs *Agrobacterium tumefaciens* to deliver viral vectors encoding viral proteins and short (∼300 bp) sequences of the target gene. A critical factor is the host range of the viral vector. Over the past years, multiple plant viruses (e.g. tobacco mosaic virus (TMV), potato virus X (PVX), and tomato golden mosaic virus (TGMV)) have been used to develop VIGS vectors. Of these, tobacco rattle virus (TRV) has the broadest host range, with results reported from numerous angiosperms (Dommes et al., 2019). To facilitate the development of VIGS methods in new species and to test the efficacy of VIGS vectors, genes of which silencing results in an easily visible phenotype are employed. Commonly used genes include *phytoene desaturase* (*PDS*), of which downregulation disrupts the conversion of phytoene to ζ-carotene, and *magnesium chelatase subunit H* (*CHL-H*), the product of which is required for the conversion of protoporphyrinogen IX to Mg-protoporphyrin IX in chlorophyll biosynthesis (Ratcliff et al., 2001; Hiriart et al., 2003). The downregulation of these genes results in white or yellow leaves (Ratcliff et al., 2001; Hiriart et al., 2003).

VIGS has been widely used in model plants such as *Nicotiana benthamiana (Kumagai et al., 1995; Hiriart et al., 2003)* and *Arabidopsis thaliana* (thale cress) (Burch-Smith et al., 2006). It has also been utilised in several crops, including *Hordeum vulgare* (barley) (Hein et al., 2005), *Oryza sativa* (rice) (Ding et al., 2007), *Zea mays* (maize) (Wang et al., 2016) and *Gossypium* (cotton) species (Tian et al., 2024). VIGS has been employed to study numerous gene families, including those involved in primary and secondary metabolism (Zulfiqar et al., 2023). Several studies have reported the use of VIGS for the discovery and characterisation of genes involved in biosynthesis of bioactives and natural products. For example, VIGS was used to discover and characterise genes involved in the production of terpenoid indole alkaloids in *Catharanthus roseus* (Madagascar periwinkle) (Liscombe and O’Connor, 2011; Yamamoto et al., 2021). It has also been used to investigate the function of genes encoding cytochrome P450s involved in the production of ursolic and oleanolic acids in *Ocimum basilicum* (sweet basil) (Misra et al., 2020). VIGS is particularly useful for studying genes involved in the biosynthesis of compounds that are essential for plant growth and development such as the role of cycloartenol synthases (CASs), of which loss-of-function early in development results in severe phenotypes or lethality (Babiychuk et al., 2008). The use of VIGS to silence *SlCAS* in *Solanum lycopersicum* (tomato) enabled the elucidation of cholesterol biogenesis (Sonawane et al., 2016). In that study, silencing of *SlCAS* led to a significant decrease in α-tomatine, cholesterol, stigmasterol, β-sitosterol, isofucosterol and cycloartenol. At the same time, the level of β-amyrin, which depends on the availability of the 2,3-oxidosqualene precursor, significantly increased in the leaves of plants in which *SlCAS*-was silenced.

The Asteraceae family is the largest and most diverse family of land plants. However, methods for gene delivery and genetic transformation are limited to a few species (Darqui et al., 2021). Stable transformation has been demonstrated in the Asteraceae crop species, *Lactuca sativa* (lettuce), *Helianthus annus* (sunflower) (Müller et al., 2001; Darqui et al., 2021; Roche et al., 2025) and the medicinal plant, *Artemisia annua* (sweet wormwood) (Alam et al., 2014; Elfahmi et al., 2014; Hassani et al., 2023). VIGS has been demonstrated in common sunflower, sweet wormwood, lettuce (Mardini et al. 2024; Wang et al. 2021; Liu et al. 2024) and has also been used to study anthocyanin biosynthesis in *Gerbera hybrida* (African daisy) (Deng et al., 2012), and chlorogenic acid and dicaffeoylquinic acid production in *Cynara cardunculus* (globe artichoke) (Moglia et al., 2016). The Asteraceae species, *Calendula officinalis* (pot marigold) has been used as a medicinal plant for thousands of years. Recently, we have shown that hydroxylated triterpenoids found in these species have potent anti-inflammatory activity (Golubova et al., 2025). To date, few studies have aimed at genetic transformation of pot marigold, however, *Rhizobium rhizogenes* has been used to produce transgenic hairy root cultures (Długosz et al., 2018; Matvieieva et al., 2025). Here, we report the development of methods for *A. tumefaciens* infiltration and VIGS for this species, applying them to silence *CAS*.

## Results

### *Agrobacterium tumefaciens*-mediated transient gene expression

We first investigated whether pot marigold is amenable to transient transfection by *A. tumefaciens*. To do this we transformed three *A. tumefaciens* strains (GV3101, LBA4404 and AGL1) with a binary vector (pEPCTαKN0001; Supplementary Table 1) encoding a chimeric gene for constitutive expression of firefly luciferase (LucF) under control of the Cauliflower Mosaic Virus 35s promoter (CaMV35sP). The resulting *A. tumefaciens* strains were infiltrated into the abaxial side of leaves of young (28 days post germination) pot marigold and *N. benthamiana* (control) plants using a 1 mL needleless syringe. Five days post-infiltration, protein was extracted and luminescence was quantified. In *N. benthamiana, A. tumefaciens* LBA4404 gave the highest levels of luminescence (Figure 1; Supplementary Table 2). Expectedly, levels of luminescence detected in pot marigold were lower than those observed in *N. benthamiana* (three orders of magnitude). Nevertheless, significant levels of luminescence were detected in plants infiltrated with *A. tumefaciens* AGL1 and *A. tumefaciens* LBA4404 (Figure 1; Supplementary Table 2). *A. tumefaciens* AGL1 gave the highest levels of expression and was used in subsequent experiments.

**Figure 1.**
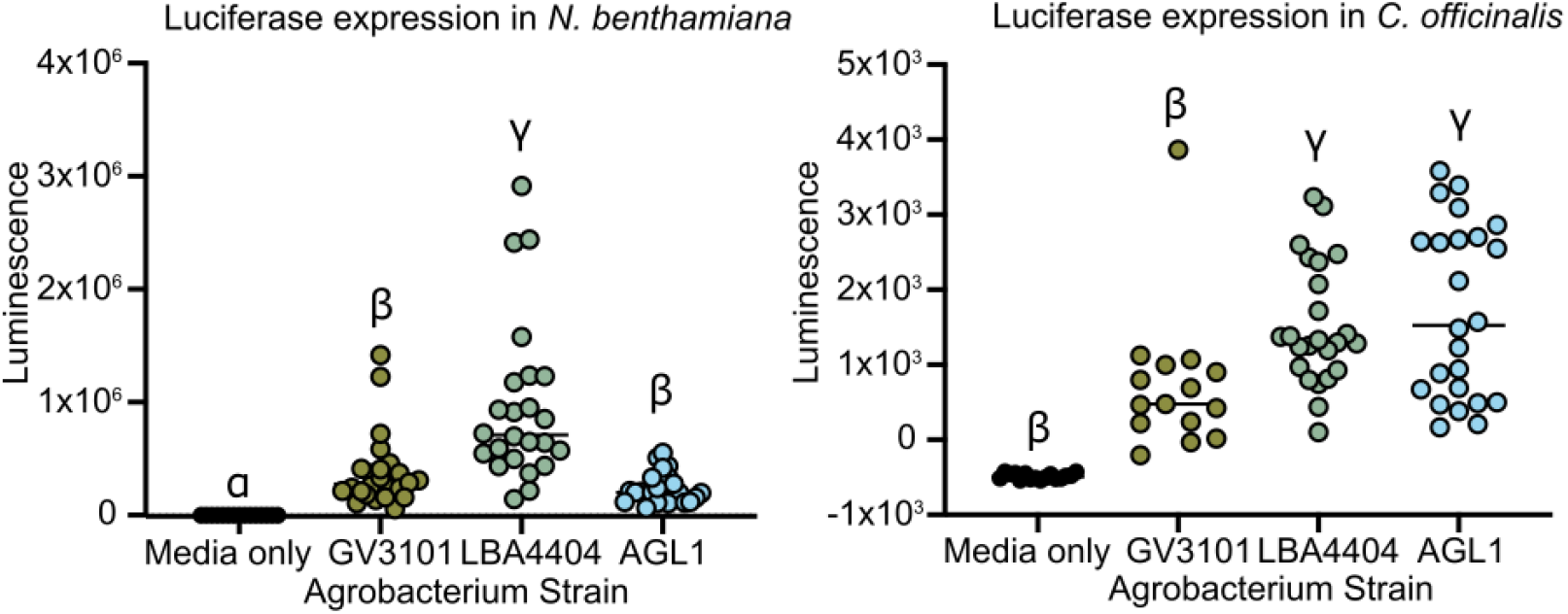
Transient expression of luciferase reporters in *Nicotiana benthamiana* and *Calendula officinalis* (pot marigold). Leaves were infiltrated with *A. tumefaciens* GV3101, LBA4404 and AGL1 carrying constructs for constitutive expression of firefly luciferase (LucF). Eight leaf disks were sampled from each of three independently infiltrated plants, except pot marigold GV3101 (two plants). Dots represent individual leaf discs. Significant differences were analysed using a Kruskal–Wallis test with a post-hoc Dunn test into groupings represented by ɑ, β or γ.

### Development of a visual marker for virus-induced gene silencing in pot marigold

Next, we identified pot marigold orthologues of *PDS* and *CHL-H* to evaluate their use as visual marker genes for VIGS. To identify candidates, previously characterised *PDS* and *CHL-H* genes from *Nicotiana* sp. (*NbPDS*; LC543532.1 and *NtCHL-H;* NM_001325713.1) were used as queries in tBLASTn to search existing pot marigold genome and transcriptome sequence data (GCA_964273985.1; PRJEB80524). Two candidate *CoPDS* genes were identified encoding proteins with 96.5 % identity to each other. Three candidate *CoCHL-H* genes were identified encoding proteins with 97.5 %-98.8 % identity to each other. The translated amino acid sequences of these genes were used to search the predicted translatomes of five Asteraceae species (*Helianthus annuus, Cynara cardunculus* var. scolymus, *Artemisia annua, Lactuca sativa* and *Cichorium endivia)* and three non-Asteraceae (*Arabidopsis thaliana, N. benthamiana* and *Amborella trichopoda*). *H. annuus, C. cardunculus* var scolymus, *L. sativa, A. thaliana, N. benthamiana PDS* genes have previously been silenced through VIGS. *A. annua, A. thaliana, N. benthamiana CHL-H* genes have also been silenced through VIGS. *A. trichopoda* was included as an outgroup as an early diverging flowering plant. *C. endivia* was included to increase the resolution of the Asteraceae. The retrieved sequences were used to construct a maximum likelihood phylogeny in which CoPDS and CoCHL candidates were observed to be most closely related to candidates from other Asteraceae (Figure 2A). To confirm which of the candidate visual selectable marker genes were expressed in our target tissues (leaves and flowers), we investigated the expression levels of *CoPDS and CoCHL* genes in three tissues of pot marigold by differential gene expression analysis of a previously published RNAseq dataset of three tissues, leaves, ray florets and disk florets (https://doi.org/10.5281/zenodo.13869958, DESeq2, Salmon v1.2.0; Golubova et al., 2025). Both *CoPDS1 and CoPDS2* were most highly expressed in pot marigold ray florets. Expression was also detected in the leaves (Figure 2B). All *CoCHL-H* genes were found to be predominantly expressed in leaves with minimal expression in floral tissues. This is expected due to its role in chlorophyll biosynthesis. *CoCHL-H2* and *3* were the most highly expressed (Figure 2B).

**Figure 2.**
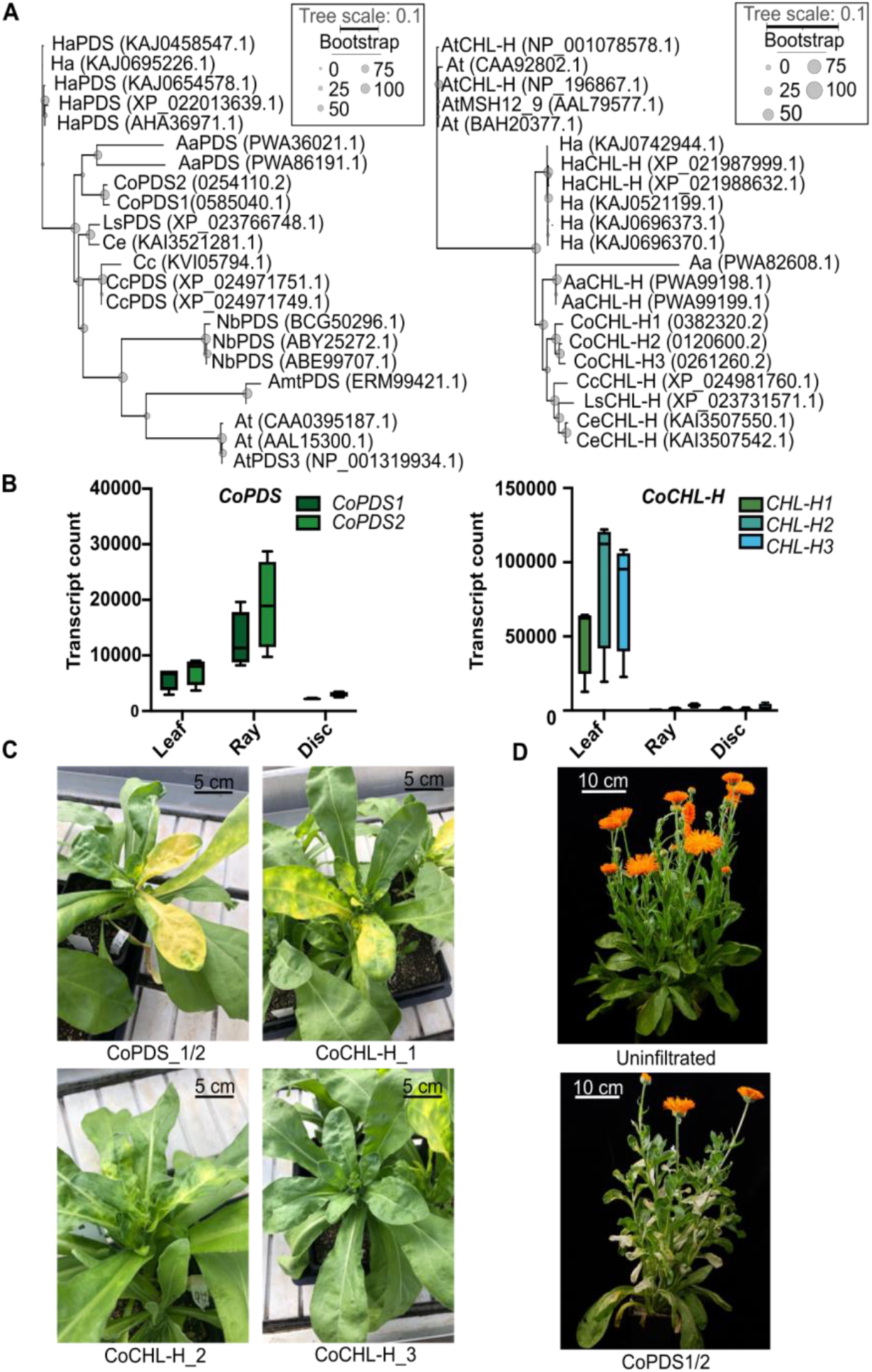
Virus-induced gene silencing of phytoene desaturase (PDS) and *magnesium chelatase subunit H* (*CHL-H*) *in Calendula officinalis* (pot marigold). (A) Maximum likelihood phylogeny of PDS and CHL-H protein sequences. *Helianthus anuus* (Ha), *Arabidopsis thaliana* (At), *Nicotiana benthamiana* (Nb), *Cynara cardunculus* var. scolymus (Cc), *Artemisia annua* (Aa), *Lactuca sativa* (Ls) and *Cichorium endivia* (Ce) and *Amborella trichopoda* (Amt). The scale bar indicates the number of substitutions. *CoPDS* and *CoCHL* genes are named according to a previously published transcriptome (https://doi.org/10.5281/zenodo.13869958) (B) Expression of candidate *CoPDS* and *CoCHL* genes in *C. officinalis* leaf, ray floret and disc floret tissues. Boxplots show normalised transcript counts of four biological replicates; the centre line denotes the median value; the boxes show the 25th to 75th percentiles and the whiskers mark the minimum and maximum values. (C) Images of representative plants infiltrated with VIGS vectors targeting *CoPDS* and *CoCHL*14 days post infiltration. Scale bar = 5 cm. (D) Images of control plant and representative plant infiltrated with VIGS vectors 35 days post infiltration. Scale bar = 10 cm.

The pTRV vectors are known to have the widest host range of established VIGS vectors and thus were selected for use. The two *CoPDS genes* share a 291 bp region in which there is only a single nucleotide polymorphism (SNP). This region was amplified and cloned into pTRV2-GG (Addgene plasmid # 105349; Supplementary Table 1). Similarly, 303 bp fragments of *CoCHL-H1, CoCHL-H2* and *CoCHL-H3* were amplified and cloned into pTRV2-GG (Addgene plasmid #105349; Supplementary Table 1). Plants were co-infiltrated at 28 days old with a strain of *A. tumefaciens* AGL1 containing pTRV1 and a strain containing pTRV2 into which fragments of the visual marker genes described above had been cloned (pTRV2-*PDS_1/2*, pTRV2-*CoCHL-H1*, pTRV2-*CHL-H_2*, pTRV2-*CoCHL-H_3*). We initially compared leaf infiltration with a 1 mL needless syringe to injection with a needle into the primary veins (midribs) of the first true leaf of 28-day-old plants (Supplementary Data 1). In plants in which leaf infiltration was used, only minimal loss of green pigment was observed around the site of infiltration, and no further analyses were performed on these plants. In plants in which pTRV-*PDS*, pTRV-*CHL-H_1* or pTRV-*CHL-H_2* was injected into the midrib, a bleaching phenotype was observed approximately 35 days post infiltration (dpi) in newly developed leaves. Surprisingly, given the comparatively low level *CoCHL-H1* expression, plants in which *CoCHL-H1* were targeted showed the most robust visual phenotype of the three *CoCHL-H* candidates (Figure 2C). However, *CoPDS* showed the most obvious phenotype and was selected for subsequent assays (Figure 2C and D).

In previous studies, infiltration with *A. tumefaciens* carrying empty a pTRV2 vector was shown to result in increased viral symptoms. In contrast, vectors in which a 300 bp fragment of a GFP coding sequence showed no symptoms (Wu et al., 2011; Broderick and Jones, 2014). Thus, an “empty vector” control VIGS vector with a visual marker was constructed by fusing the previously tested fragment of *CoPDS* to a 300 bp fragment of GFP, creating pTRV2(PDS:GFP). The efficacy of this vector was tested by co-infiltration with pTRV1 into plants as previously described. Thirty-eight days post infiltration, RNA was extracted from leaf tissues in which the leaves showed a bleaching phenotype and the expression level of two reference genes (*SAND* and *PROTEIN PHOSPHATASE 2A (PP2A)*) and *CoPDS1 and CoPDS2* were assessed by qRT-PCR. Infiltration with pTRV1+pTRV2(PDS:GFP) reduced the expression of *CoPDS1* (92.1%) and *CoPDS2* (91.5%) (Supplementary Figure 1; Supplementary Table 3).

### Virus-induced gene silencing of cycloartenol synthases

Five putative cycloartenol synthases were previously identified in pot marigold but had not been functionally characterised (Golubova et al., 2025). The protein sequences of these putative cycloartenol synthases (CoCAS1, 2, 3, 4 and 5) were used as a queries in tBLASTn to search genomes of *Helianthus anuus* (Ha), *Arabidopsis thaliana* (At), *Nicotiana benthamiana* (Nb), *Cynara cardunculus* var. scolymus (Cc), *Artemisia annua* (Aa), *Lactuca sativa* (Ls), *Cichorium endivia* (Ce) and *Amborella trichopoda* (Amt). Protein sequences with at least 87% coverage and 65.7% identity were selected as candidate CASs, aligned and used to contrast a maximum likelihood phylogenetic tree (Figure 3A). The relative gene expression of the *CoCAS* orthologues across leaf and floral tissues was analysed as described above and *CoCAS2* and *4* were found to be highly expressed in leaves and selected as targets for VIGS (Figure 3B). To construct VIGS vectors for *CoCAS2* and *4*, a 300 bp region in which sequence identity of these two genes is 100% was identified. This region was amplified and cloned into pTRV2 fused to *CoPDS* (Figure 3C; Supplementary Table 1). Subsequently, plants were co-infiltrated with pTRV1 and either the control plasmid (pTRV2-PDS:GFP) or the experimental plasmid (pTRV2-PDS:CAS).

**Figure 3.**
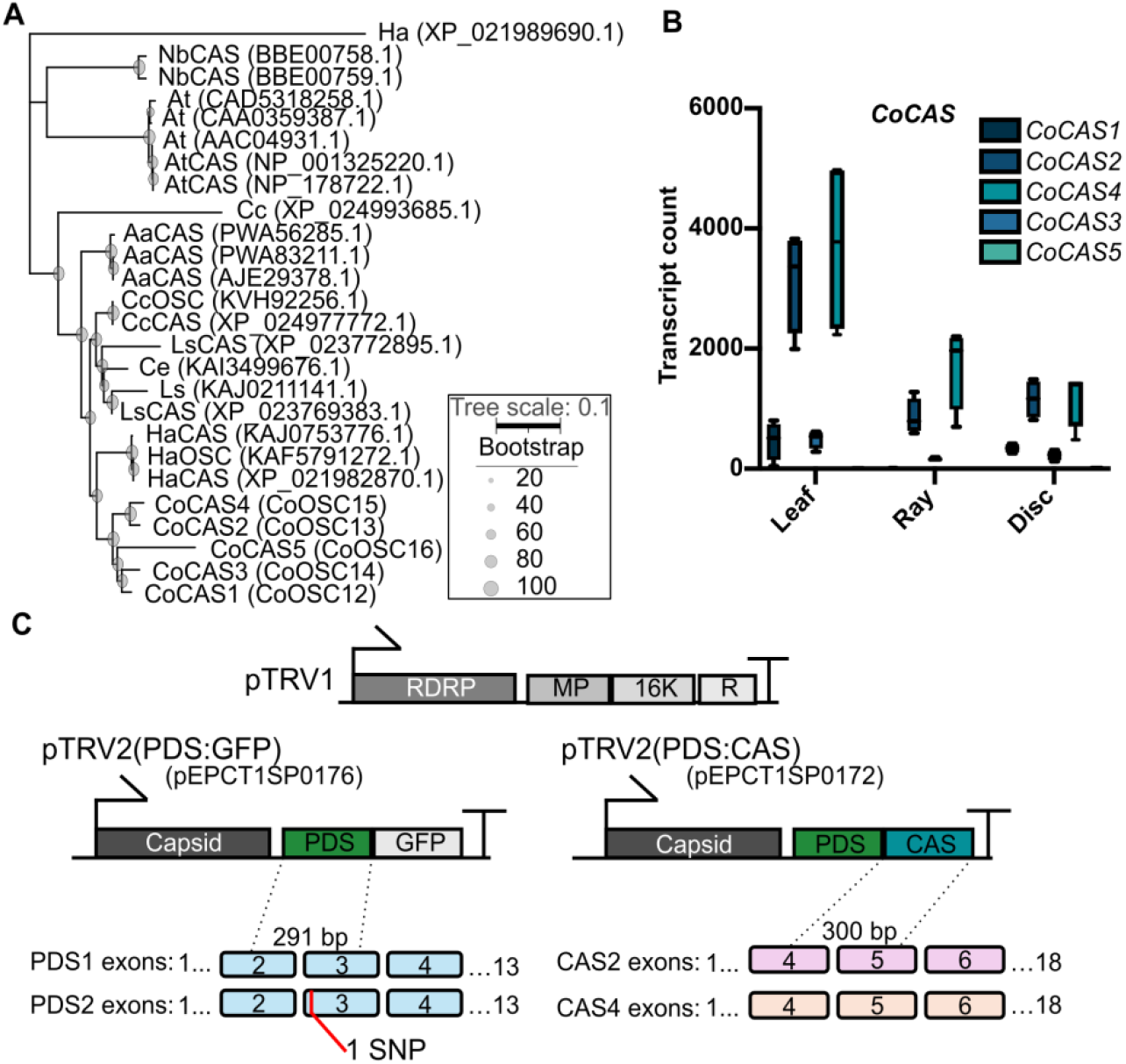
Identification of candidate *Calendula officinalis* (pot marigold) *cycloartenol synthase (CAS)* genes. (A) Maximum likelihood phylogeny of *CASs*. Scale bar indicates the number of substitutions. *Helianthus anuus (Ha), Arabidopsis thaliana (At), Nicotiana benthamiana (Nb), Cynara cardunculus var. scolymus (Cc), Artemisia annua (Aa), Lactuca sativa (Ls) and Cichorium endivia (Ce), Amborella trichopoda (Amt)*. CoCAS 1-5 were identified in Golubova *et al*., 2025. The names in parentheses (OSC) are those assigned in that publication. (B) Expression of candidate *CoCAS* genes in *C. officinalis* leaf, ray floret and disc floret tissues. Boxplots show normalised transcript counts of four biological replicates; the centre line denotes the median value; the boxes show the 25th to 75th percentiles and the whiskers mark the minimum and maximum values. (C) Schematics of plasmids for virus-induced gene silencing. A fragment of the visual marker gene *CoPDS1/2* is fused to a fragment of either a control (*GFP*) or target gene (*CAS*).

### Reduced expression of pot marigold cycloartenol synthases alters sterol accumulation

The first committed step of phytosterol biosynthesis is the conversion of 2,3 oxidosqualene to cycloartenol by CASs. This is then converted to campestrol (used to produce brassinolides), or to stigmasterol via isofucosterol and β-sitosterol, which are all integral membrane components (Figure 4A). First, the expression levels of the target genes were analysed. Samples were taken from leaves that showed a bleaching phenotype 38 days post infiltration. RNA was extracted and gene expression analysed by qRT-PCR. The expression levels of *CoCAS2* and *CoCAS4* were significantly reduced (69.0% and 70.5% respectively) in plants infiltrated with pTRV1+ pTRV-PDS:CAS compared to the control (Figure 4B; Supplementary Table 4). Samples from the same leaves were extracted in ethyl acetate, dried and derivatised with N-Methyl-N-(trimethylsilyl)trifluoroacetamide and analysed by gas chromatography–mass spectrometry (GC–MS). Peaks were identified by comparison to authentic standards (Supplementary Figure 2). Levels of campesterol and β-sitosterol were comparable between those infiltrated with the control plasmid (pTRV2-PDS:GFP) and those targeting *CoCAS2* and *4* (pTRV2-PDS:CAS) (Figure 4C; Supplementary Table 4). However, the accumulation of stigmasterol was reduced in plants in which *CoCAS2* and *4* expression was reduced and the accumulation of isofucosterol was increased (Figure 4C; Supplementary Table 4). To confirm that silencing of *CoPDS1/2* did not affect sterol accumulation, sterol levels were also compared to uninfiltrated plants, and no significant differences were observed (Supplementary Figure 3; Supplementary Table 5).

**Figure 4.**
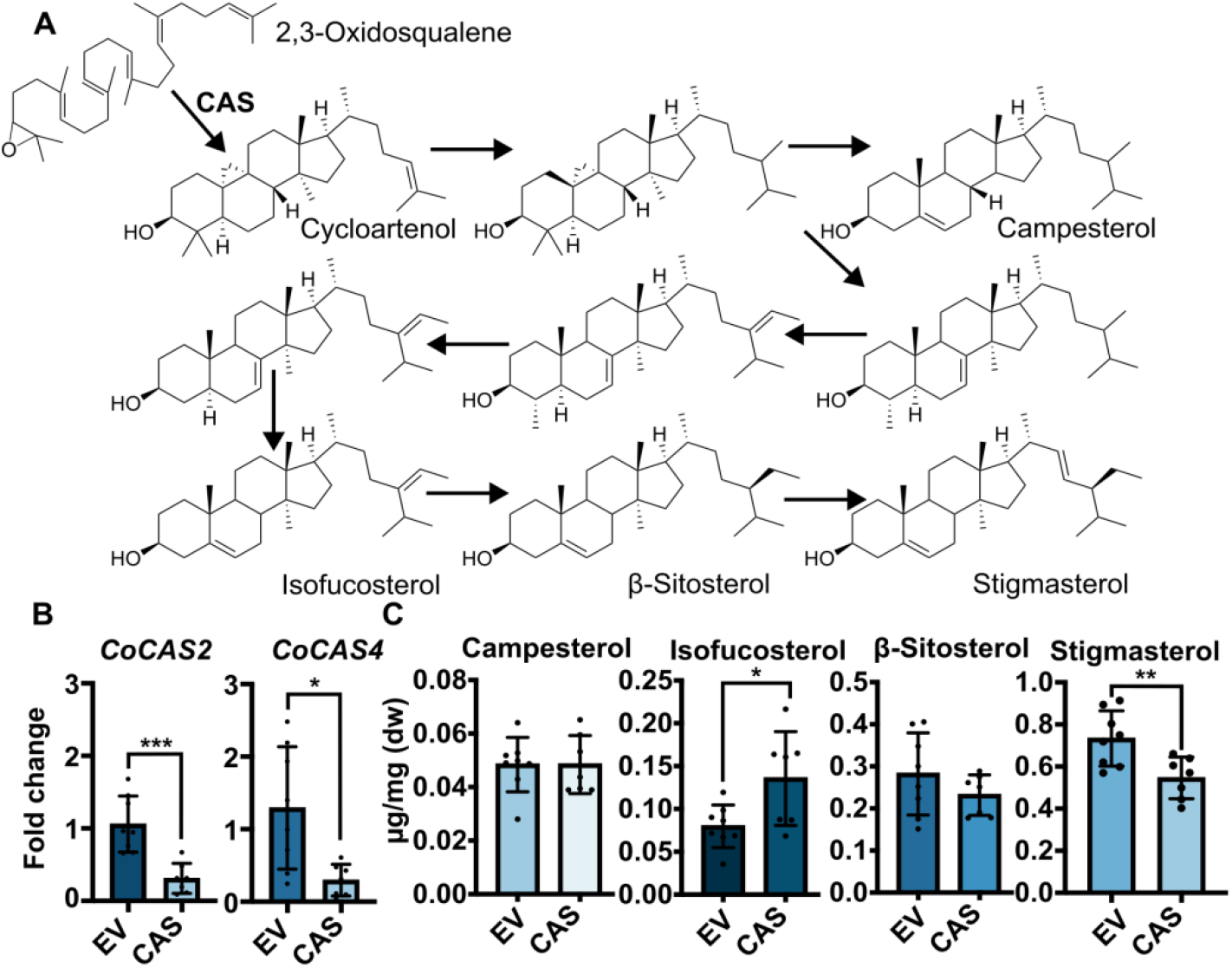
Virus induced gene silencing of *cycloartenol synthases* reduced the accumulation of stigmasterol. (A) Schematic of the conversion of 2,3-oxidosqualene to cycloartenol by CAS and subsequent modifications to campesterol, isofucosterol, β-sitosterol and stigmasterol (B) Expression levels of *CoCAS* genes in pot marigold leaves 38 days post infiltration with VIGS vectors as determined by qRT-PCR; n ≥ 7. Bars show the mean and standard error of four replicates. Statistical significance was determined for *CoCAS2* using a Student’s t-test. For *CoCAS4*, a Mann-Whitney U test was used as there was no evidence of equal variance or normal distribution (Supplementary Table 4). (C) Quantification of stigmasterol, isofucosterol, β-sitosterol and campesterol in plants in which expression of *CoCAS* was reduced by VIGS. Metabolites were quantified by GC-MS using an internal standard. n ≥ 7; statistical significance was calculated using a Student’s t-test (Supplementary Table 4); EV = empty vector (pTRV2-PDS:GFP); CAS= pTRV2-PDS:CAS.

Finally, to assess if VIGS can be used to study florally expressed genes, samples of ray floret tissues were collected from the flowers of plants infiltrated with the same constructs 38 days post-infiltration. In addition, a second set of plasmid vectors was constructed in which a fragment of *FLOWERING LOCUS T*, previously reported to facilitate transport of infiltrated small RNAs to floral meristems (Ellison et al., 2020), was fused to the 3’ end of the PDS and CAS gene fragments (Supplementary Table 1). However, no reduction in the expression of the target genes was identified in the flowers using either set of vectors (Supplementary Figure 4; Supplementary Table 6).

## DISCUSSION

VIGS is a powerful method for functional characterisation of endogenous plant genes. To establish a VIGS method for pot marigold, we first investigated agroinfiltration, finding that *A. tumefaciens* AGL1 and LBA4404 gave the highest levels of expression (Figure 1). *A. tumefaciens* AGL1 has a similar chromosomal background to the wild-type strain, C58, containing a 2.8 Mb circular chromosome and a 2 Mb linear chromosome (De Saeger et al., 2021). It also contains the pTiBo542 tumour-inducing (Ti) plasmid, sometimes called a hypervirulence plasmid due to the presence of additional copies of the *virulence* (*vir*) genes, and a 0.54 Mb pAtC58 accessory plasmid. Transformation efficiency has been shown to increase with *A. tumefaciens* strains containing pTiBo542 (Komari, 1990). Further, AGL1 carries an insertion mutation in its *recA* general recombination gene (Lazo et al., 1991), which may also impact plasmid integrity and therefore efficiency of transformation. *A. tumefaciens* LBA4404 has an Ach5 chromosomal background, which is also similar to C58, (De Saeger et al., 2021). Both AGL1 and LBA4404 are widely used and have been shown to transform numerous plant species (Lee and Gelvin, 2008).

In this study, VIGS was achieved by injecting *A. tumefaciens* into the midrib vein of immature (non-flowering) pot marigold plants (Figure 4; Supplementary Data 1). The use of *PDS1* as a marker gene enabled viral spread to be tracked, guiding the sampling of tissues (Figure 1; Figure 4). This was important as silencing only occurred in some sectors of some leaves. Alternative viral vectors may improve infection as might different methods for delivery of *A. tumefaciens*. Previously, vacuum infiltration of seeds improved VIGS efficiency in sunflowers. In *Nepeta* sp. (catmint), *A. tumefaciens* dropped onto wounds enabled VIGS and in sweet wormwood the use of the surfactants Silwet L-77 and Silwet S-408 improved the efficiency of *A. tumefaciens*-mediated transient gene expression (Elfahmi et al., 2014; Palmer and O’Connor, 2020; Mardini et al., 2024).

Several secondary metabolites are known to be controlled via feedback mechanisms (Li et al., 2024a). VIGS of *CoCAS2* and *CoCAS4* resulted in less stigmasterol but not of campesterol, isofucosterol, or β-sitosterol (2024). Although more detailed investigations are required, this suggests mechanisms exist to maintain some membrane components and flux to brassinolides, possibly by feedback at the protein or the transcriptional level. Silencing of *CAS* genes has been observed to have different effects in other species: In tomato, silencing of *SlCAS* reduced the accumulation of cholesterol, stigmasterol, *β*-sitosterol, isofucosterol and cycloartenol, revealing that SlCAS catalyses the production of common precursor for tomato cholesterol and phytosterol metabolism (Sonawane et al., 2016). In *N. benthamiana*, silencing *NbCAS1* increased the expression of *NbHMGR1*, which acts further up in the mevalonate pathway converting 3-hydroxy-3-methyl 3-glutaryl coenzyme A to mevalonate, which is used to produce the 2,3-oxidosqualene precursor for sterol production. In contrast, expression of *NbSSR2* and *NbSMT1*, which encode enzymes involved in the production of steroidal glycoalkaloids and sterols respectively, were reduced (Atsumi et al., 2018). These studies demonstrate that VIGS can be combined with quantification of transcripts and metabolites to investigate the regulation of metabolic pathways. In the future, comparative studies may reveal commonalities and divergences in sterol metabolism between plant lineages. Demonstrating efficient VIGS methods for new species contributes to this aim.

There are few reports of VIGS in floral tissues, though TRV-based vectors have been used to study flower floral senescence in *Petunia hybrida* (Chen et al., 2004). TRV is known to systemically infect this and other solanaceae species. VIGS of floral tissues was also demonstrated in *Rosa chinensis* (China rose) and used to study anthocyanin biosynthesis (Yan et al., 2020). In Asteraceae, VIGS and overexpression constructs were infiltrated into the detached petals of *Gerbera hybrida* (Jiang et al., 2022), but entry of viruses into floral meristems is limited (Li et al, 2024b). Some plant mRNAs are mobile and can move between cells or organs to transmit environmental signals into developmental programs (Luo et al., 2024). For example, the mRNA of *FLOWERING TIME* (*FT*), which encodes for a mobile protein that mediates the onset of flowering, was itself shown to move from leaves to the shoot apical meristem (SAM) through the phloem (Pin and Nilsson, 2012; Liu et al., 2013). In 2020, Ellison et al. reported that TRV vectors expressing single guide RNAs (sgRNA) could be used to produce heritable mutations following transient expression in the leaves of *N. benthamina* plants constitutively expressing Cas9 (Ellison et al., 2020). They reported that the addition of a *FT* tag to sgRNAs increased the frequency of heritable edits by promoting entry into the shoot apical meristem, which has been successfully employed to edit targeted in several studies including our own (Dudley et al., 2022; Vollheyde et al., 2023). In this study, we fused an identical *FT* tag to target RNAs in VIGS constructs. However, this did not enable gene silencing in floral tissues of pot marigold. The identification of alternative VIGS vectors with systemic infection may improve VIGS in floral tissues of Asteraceae.

## METHODS

### Plant growth

Seeds of *C. officinalis* (pot marigold) were purchased from Chiltern Seeds (Wallingford, UK). Seeds were sown in 9 cm plastic pots with Levington F2 starter (100% peat). Approximately ten days post-emergence, seedlings were transplanted into 11 cm plastic pots containing 60% peat, 20% grit, 20% perlite, 2.25 kg/m^3^ dolomitic limestone, 1.3 kg/m^3^ PG mix, 3 kg/m^3^ osmocote exact. All plants were grown in summer glasshouse conditions with natural day length and temperature. *N. benthamiana* was sewn in a 10 cm plastic pot and cultivated in a peat-based potting mix (90% peat, 10% grit, with 4 kg/m3 dolomitic limestone, 0.75 kg/m3 powdered compound fertiliser, 1.5 kg/m3 slow-release fertiliser). *N. benthamiana* plants were grown in a controlled environment room with 16 h light, 8 h dark at 22 °C, 80% humidity and ∼200 µmol/m2/s light intensity for 28 days prior to infiltration.

### Transformation of *A. tumefaciens*

Aliquots of 40 μL electrocompetent *A. tumefaciens* were incubated on ice with 200 ng of plasmid DNA for 2 min. Electroporation was performed in ice-chilled 2 mm electroporation cuvettes (Geneflow) using a MicroPulser Electroporator (Bio-Rad) at 2.2 kV with <1 s pulse duration. Immediately after electroporation, 400 μL of LB were added and cells were transferred to 2 mL microcentrifuge tubes and incubated at 28 °C for 2-3 h at 200 rpm. 20 μL was plated onto LB-agar with appropriate antibiotics, using 6 mm glass beads. Plates were dried for 5 min and incubated upside down at 28 °C for 48 h.

### Transient expression of luciferase

*A. tumefaciens* strains AGL1, LBA4404 and GV3101 harbouring the firefly luciferase (LucF) under the control of the Cauliflower Mosaic Virus 35 s promoter (CaMVp) (pEPCTɑKN0001; Addgene #187568) were grown in LB medium supplemented with 50 μg/mL kanamycin plus ether 100 μg/mL rifampicin and 50 μg/m carbenicillin (AGL1); 100 μg/mL rifampicin and 50 μg/m streptomycin (LBA4404) or 100 μg/mL rifampicin and 20 μg/mL gentamicin (GV3101) respectively for each *A. tumefaciens* strain. Details of all plasmid vectors are provided in Supplementary Table 1 and details of *A. tumefaciens* strains are provided in Supplementary Table 7. Cultures were incubated for 16 h at 28 °C/220 rpm. Overnight saturated cultures were centrifuged at 3000 *g* for 20 min at room temperature, and cells were resuspended in infiltration medium (10 mM 2-(N-morpholino)ethanesulfonic acid (MES) pH 5.6, 10 mM MgCl2, 200 μM 3′,5′-dimethoxy-4′-hydroxyacetophenone (acetosyringone)) to OD600 0.8, and incubated at room temperature for 2-3 hrs. *A. tumefaciens* cultures were infiltrated into the abaxial side of leaves of 28 day-old non-flowering *N. benthamiana* and pot marigold plants with 3-4 fully expanded true leaves using a 1 mL needleless syringe. Plants were grown in controlled environment chambers (16 h light, 8 h hours dark; 22 °C; 120– 180 μmol/m2/s light intensity). After five days, two 1 cm diameter discs were sampled from infiltrated leaves. Luciferase expression was detected using the Luciferase® reporter assay system (Promega, Madison, WI). Leaf tissue was homogenised in 180 μL passive lysis buffer (Promega) with protease inhibitor (P9599, Sigma-Aldrich, Dorset, UK). Following incubation on ice for 15 min and centrifugation (14 000 *g*; 2 min; 4 °C), the supernatant was diluted 1:10 in passive lysis buffer. 10 μL of this dilution was further diluted 1:3 in passive lysis buffer and an equal volume (30 μL) of ONE-Glo™ EX Luciferase Assay Reagent (Promega) was added. After 10 min at room temperature, luminescence was quantified using a GloMax 96 Microplate Luminometer (Promega) with a 10 s read time and 1 s settling time.

### Identification of candidate genes and phylogenetic analysis

Candidate *CoPDS* and *CoCHL-H* genes were identified using tBLASTn to search the pot marigold genome using NbPDS (LC543532.1) and NtCHL-H (NM_001325713.1) as queries. The identification of *CoCAS* was previously described (Golubova et al., 2025). The translated protein sequences of the retrieved pot marigold genes were used to search the publicly available translatomes of *A. trichopoda* (txid:13333), *A. thaliana* (txid:3702), *N. benthamiana* (txid:4100), *A. annua* (txid:35608), *L*.*sativa* (txid:4236), *C. endive* (txid:114280), *T. kok-saghyz* (txid:333970), *H*.*anuus* (txid:4232), *C*.*seticuspe* (txid:1111766), *C. cardunculus* (txid:4265). Sequences with 87% coverage and 65.7% identity to CoCAS, 77% coverage and protein identity to CoCHL-H and 91% coverage and 81.7% identity to CoPDS candidates were selected. Protein sequences were aligned using MAFFT (v7.511) and trimmed using ClipKIT with smartgap mode (v2.1.3). Maximum likelihood trees were inferred using IQ-TREE and models were selected using ModelFinder (CAS=JTTDCMut+G4, CHL-H=VT+G4, PDS =JTT+i+G4) (Kalyaanamoorthy et al., 2017). Differential expression analysis was performed as previously described using existing data (https://doi.org/10.5281/zenodo.13869958) (Golubova et al., 2025).

### Construction of plasmid vectors for virus induced gene silencing

Target sequences were amplified from cDNA or existing plasmids using primers with extension that enabled cloning into BsaI sites of pTRV2 (Addgene #105349). Primer sequences are provided in Supplementary Table 8. Constructs were assembled using a one-step digestion-ligation reaction as previously described (Cai et al., 2020). Sequence verified constructs were transformed into *A. tumefaciens* AGL1 by electroporation. The details of all plasmid vectors are provided in Supplementary Table 1.

### *Agrobacterium* infiltration for VIGS

Cultures of electrocompetent *A. tumefaciens* AGL1 with pTRV1 (pNJB069) or pTRV2 (Addgene #105349) were grown as described above. Cells were collected by centrifugation and resuspended in infiltration medium with acetosyringone to OD_600_ 0.8. Cultures with pTRV1 and pTRV2 were in a 1:1 ratio prior to infiltration. For *N. benthamiana*, 28 days old plants were infiltrated as described above. The first true leaves of 28-day-old pot marigold plants were infiltrated as described above or by using a 1 mL syringe with a needle to inject cultures into the midrib until liquid was observed to spread through the plant (Supplementary Data 1).

### Purification of nucleic acids

Two samples of total RNA were purified from leaf and ray floret tissue collected from four independent plants per construct, snap-frozen frozen in liquid nitrogen and stored at -70 °C. RNA was extracted using the Spectrum Plant Total RNA Kit (STRN250; Merck, Darmstadt, Germany) following the manufacturer’s instructions except that during column washing, 300 μL of column wash 1 was used to wash RNA and then 80 μL of DNase I (RNA-Free Dnase Set 79254; Qiagen, Hilden, Germany) was added to the column and incubated for 15 minutes followed by 500 μL of wash 1. RNA integrity was assessed by electrophoresis on a 1.5 % agarose gel and using a Bioanalyzer (Agilent Technologies, Sata Clara, CA, USA).

### Quantitative reverse transcription PCR (qRT-PCR)

Reverse transcription was perfumed using M-MLV cDNA (Sigma: M1302) according to the manufacturer’s instructions. Following first strand synthesis, reactions were inactivated at 70 °C for 15 minutes and cDNA was diluted 1:10 in TE (5 mM Tris-HCl + 0.5 mM EDTA, pH 8.0). As previously reported, SAND and PP2A were used as reference genes (Golubova et al., 2025). Amplification was performed in 10 μL reactions (0.2 μM each primer, 6 ng cDNA, 1 μL SYBR Green Jumpstart™ Taq ReadyMix™) and incubated at 94 °C for 2 minutes and 40 x (94 °C for 15 seconds and 58 °C for 60 seconds). Two qPCR reactions (technical replicates) were performed for each of four biological samples (independently infiltrated plants). Control reactions (no reverse transcriptase; no template controls) had Cq values of >35. Relative expression analysis was calculated using the ΔΔCt method.

### Metabolite extraction and GC-MS analysis

Metabolites were extracted from 30 mg of freeze-dried plant material by homogenisation with Samples were mixed in 500 μL ethyl acetate (Sigma Aldrich, Burlington, MA, USA) at 700 rpm at 40 °C for 2 hours incubated at room temperature for 48 hours. Following centrifugation at 21,130 x *g* for 5 minutes, 50 μL of supernatant was transferred to a 2 mL glass vial and dried in a centrifugal evaporator. Samples were derivatised with 50 μL N-Methyl-N-(trimethylsilyl)trifluoroacetamide (Sigma Aldrich) at 37 °C for 30 minutes and transferred to glass inserts in 2 mL glass vials for GC-MS analysis on a 7890B GC (Agilent; Santa Clara, CA, USA) with a Zebron ZB5-HT Inferno column (Phenomenex; Washington D.C, USA) as previously described (Golubova et al., 2025). Data analysis was carried out using MassHunter workstation software (Agilent).

### Statistics and reproducibility

Analyses including metabolomics, transcriptomics and gene expression via qPCR were performed with a minimum of 6 biological replicates. Outliers were identified using a Grubbs, which was used to exclude two replicates from uninfiltrated plants and a single replicate from pTRV2(PDS:CAS) (Supplementary Table 9). The investigators were not blinded during experiments or analysis. Exact P-values and test statistics are provided in Supplementary Tables 2-6 alongside tested assumptions for each statistical test.

## Supporting information

Combined Supplementary Figures and Tables

## Author Contributions

CT, DG and NJP conceptualised the study. DG performed luciferase assays. MS, DG and CT identified candidate genes and performed phylogenetic analyses. MS, DG and CT extracted RNA. DG and CT cloned pTRV plasmids. CT and DG performed qRT-PCR. CT performed metabolic profiling. MS, CT, and NJP were responsible for supervision and project management. NJP was responsible for fundraising. DG, CT, and NJP drafted the figures and text. All authors contributed to revisions and editing.

## Acknowledgements

Plasmids pNJB069 (TRV1) was a gift from Daniel Voytas, and pTRV2 (Addgene #105349) was a gift from Johannes Stuttmann. We also thank the John Innes Centre horticultural services team for assistance with plant husbandry and the John Innes Centre metabolomics facility for assistance with GC-MS. Thanks to Sasha Stanbridge from the Earlham Institute for assistance with photography of *C. officinalis* plants. Reference standards: β-sitosterol, stigmasterol and campesterol were kindly provided by Anne Osbourn, John Innes Centre, Norwich, UK.

