## Supplementary material for "Virus Induced Gene Silencing in *Calendula officinalis* (pot marigold)": Combined Supplementary Figures and Tables

***The following supporting information is available for this article:***

**Tables**

Supplementary Table 1: Table of Plasmids
Supplementary Table 2: Table of Statistics for Figure 1
Supplementary Table 3: Table of Statistics for Supplementary Figure 1
Supplementary Table 4: Table of Statistics for Figure 4
Supplementary Table 5: Table of Statistics for Supplementary Figure 3
Supplementary Table 6: Table of Statistics for Supplementary Figure 4
Supplementary Table 7: Table of agrobacterium strains
Supplementary Table 8: Table of Primers
Supplementary Table 9: Table of Statistics for outliers

**Figures**

Supplementary Figure 1: Expression of visual markers
Supplementary Figure 2: GCMS standards and mass spectra for sterols
Supplementary Figure 3: Sterol content in controls
Supplementary FIgure 4: VIGS in floral tissues

**Data**

Supplementary Data 1: Video showing injection of *A. tumefaciens* cultures into midrib veins of *C. officinali*s

**Supplementary Table 1. List of plasmids.**

| Plasmid ID | Addgene Code | Description | Antibiotic Resistance | Source |
|---|---|---|---|---|
| pEPCTαKN0001 | 187568 | p35S:LucF:t35S | Kanamycin | Kallam et al (2023); doi: 10.1111/pbi.14048 |
| pTRV2-GG | 105349 | VIGS acceptor plasmid | Spectinomycin | Johannes Stuttmann; Ganter et al; doi: 10.1371/journal.pone.0197185. |
| pTRV1 (pNJB069) | N/A | VIGS helper plasmid | Kanamycin | Daniel Voytas; Ellison et al; doi: 10.1038/s41477-020-0670-y |
| pEPCT1SP0031 | 250239 | pTRV2:CoCHL-H.1 | Spectinomycin | This study |
| pEPCT1SP0032 | 250240 | pTRV2:CoCHL-H.2 | Spectinomycin | This study |
| pEPCT1SP0033 | 250241 | pTRV2:CoCHL-H.3 | Spectinomycin | This study |
| pEPCT1SP0034 | 250242 | pTRV2:CoPDS | Spectinomycin | This study |
| pEPCT1SP0172 | 250243 | pTRV2:CoPDS:CoCAS | Spectinomycin | This study |
| pEPCT1SP0173 | 250244 | pTRV2:CoPDS:GFP | Spectinomycin | This study |
| pEPCT1SP0175 | 250245 | pTRV2:CoPDS:GFP_FT | Spectinomycin | This study |

**Supplementary Table 2. Table of statistics for Figure 1**.
*=P<0.05, ** P<0.01,*** P<0.0001

| Experiment | Test | Sample | Value | P-Value |
|---|---|---|---|---|
| Luciferase expression in *N. benthamiana* | Levene's test | Negative, GV3101, AGL1, LBA4404 | F=9.3935 | 0.0003518 (***) |
| | Shapiro-Wilk test | Negative, GV3101, AGL1, LBA4404 | W=0.71421 | 0.000000007388 (***) |
| | Kruskal-Wallis | Negative, GV3101, AGL1, LBA4404 | $X^2$=46.223 | 0.0000000005086 (***) |
| | Dunns test | AGL1-GV3101 | Z=-1.006421 | 0.3412130 (n.s.) |
| | | AGL1-LBA4404 | Z=-3.209155 | 0.001996888 (**) |
| | | GV3101-LBA4404 | Z=-2.202733 | 0.03313626 (*) |
| | | AGL1-Negative | Z=3.456013 | 0.001096459 (**) |
| | | GV3101-Negative | Z=4.462434 | 0.00002431018 (***) |
| | | LBA4404-Negative | Z=6.665167 | 0.0000000001586185 (***) |
| Luciferase expression in *C. officinalis* | Levene's test | Negative, GV3101, AGL1, LBA4404 | F=0.8909 | 0.0.4494 (***) |
| | Shapiro-Wilk test | Negative, GV3101, AGL1, LBA4404 | W=0.12019 | <2.2e-16 (***) |
| | Kruskal-Wallis | Negative, GV3101, AGL1, LBA4404 | $X^2$=51.725 | 0.000000000003428 (***) |
| | Dunns test | AGL1-GV3101 | Z=3.21474057 | 0.001958437 (**) |
| | | AGL1-LBA4404 | Z=0.05084827 | 0.9594464 (n.s) |
| | | GV3101-LBA4404 | Z=-3.16389230 | 0.00186093 (**) |
| | | AGL1-Negative | Z=6.26361693 | 0.000000002256893 (***) |
| | | GV3101-Negative | Z=3.38826555 | 0.001406723 (**) |
| | | LBA4404-Negative | Z=6.21813686 | 0.0000000001509278 (***) |

**Supplementary Table 3. Table of Statistics for Supplementary Figure 1.**
*=P<0.05, ** P<0.01,*** P<0.0001

| Experiment | Test | Sample | Value | P-Value |
|---|---|---|---|---|
| *PDS1* expression in leaf for WT, PDS_GFP and PDS_CAS | Lavene's test | WT, PDS_GFP and PDS_CAS | F=2.8841 | 0.08057 (n.s.) |
| | Shapiro-wilk test | WT, PDS_GFP and PDS_CAS | W=0.71452 | 0.00003011 (***) |
| | Kruskal-Wallis test | WT, PDS_GFP and PDS_CAS | $X^2$=14.464 | 0.000723 (***) |
| | Dunn test | PDS_CAS – PDS-GFP | Z=-0.8767175 | 0.3806401098 (n.s.) |
| | | PDS_CAS-WT | Z=-3.6218797 | 0.0008774105 (***) |
| | | PDS_GFP-WT | Z=-2.8639438 | 0.0062760342 (**) |
| *PDS2* expression in leaf for WT, PDS_GFP and PDS_CAS | Lavene's test | WT, PDS_GFP and PDS_CAS | F=7.73 | 0.003497 (**) |
| | Shapiro Wilk test | WT, PDS_GFP and PDS_CAS | W=0.73222 | 0.00005177 (***) |
| | Kruskal Wallis test | WT, PDS_GFP and PDS_CAS | $X^2$=14.007 | 0.0009087 (***) |
| | Dunn test | PDS_CAS – PDS-GFP | Z=-0.5579111 | 0.576905075 (n.s.) |
| | | PDS_CAS-WT | Z=3.4572488 | 0.001637162 (**) |
| | | PDS_GFP-WT | Z=-3.0127201 | 0.003883764 (**) |

**Supplementary Table 4. Table of statistics for Figure 4.**
*=P<0.05, ** P<0.01,*** P<0.0001

| Experiment | Test | Sample | Value | P-Value |
|---|---|---|---|---|
| *CAS2* expression in leaf for PDS_GFP and PDS_CAS | Lavene's test | PDS_GFP and PDS_CAS | F=1.7593 | 0.2075 (n.s.) |
| | Shapiro Wilk test | PDS_GFP and PDS_CAS | W=0.93875 | 0.3667 (n.s) |
| | T-Test | PDS_GFP and PDS_CAS | T=4.3563, df=13 | 0.0007779 (***) |
| *CAS4* expression in leaf for PDS_GFP and PDS_CAS | Lavene's test | PDS_GFP and PDS_CAS | F=10.861 | 0.005796 (**) |
| | Shapiro Wilk test | PDS_GFP and PDS_CAS | W=0.85247 | 0.01882 (*) |
| | Mann-Whitney U test | PDS_GFP and PDS_CAS | W=49 | 0.01399 (*) |
| **Experiment** | **Test** | **Sample** | **Value** | **P-Value** |
| Leaf Stigmasterol content for PDS_GFP and PDS_CAS | Lavene's test | PDS_GFP and PDS_CAS | F=0.7115 | 0.4142 (n.s.) |
| | Shapiro Wilks test | PDS_GFP and PDS_CAS | W=0.96511 | 0.7802 (n.s.) |
| | T-test | PDS_GFP and PDS_CAS | T=3.0686, df= 13 | 0.008973 (***) |
| Leaf Campesterol Content for PDS_GFP, PDS_CAS | Lavenes test | PDS_GFP, PDS_CAS | F=0.3761 | 0.5503 (n.s.) |
| | Shapiro Wilks test | PDS_GFP, PDS_CAS | W=0.96844 | 0.8343 (n.s.) |
| | T-test | PDS_GFP, PDS_CAS | T=-0.0040762, df=13 | 0.9968 (*) |
| Leaf Isofucosterol Content for PDS_GFP, PDS_CAS | Lavene's test | PDS_GFP, PDS_CAS | F=2.6394 | 0.1282 (n.s.) |
| | Shapiro Wilks test | PDS_GFP, PDS_CAS | W=0.89208 | 0.07211 (n.s.) |
| | T-test | PDS_GFP, PDS_CAS | T=-2.6014, df=13 | 0.02195 (*) |
| Leaf β-Sitosterol Content for PDS_GFP, PDS_CAS | Lavene's test | PDS_GFP, PDS_CAS | F=4.329 | 0.05781 (n.s.) |
| | Shapiro Wilks test | PDS_GFP, PDS_CAS | W=0.92726 | 0.2482 (n.s) |
| | T-test | PDS_GFP, PDS_CAS | T=0.2382, df=13 | 0.2382 (n.s) |

**Supplementary Table 5. Table of Statistics for Supplementary Figure 3**.
*=P<0.05, ** P<0.01,*** P<0.0001

| Experiment | Test | Sample | Value | P-Value |
|---|---|---|---|---|
| Campesterol | Lavene's test | WT, PDS_GFP | F=0.0369 | 0.8503 (n.s) |
| | Shapiro-Wilks test | WT, PDS_GFP | W=0.92762 | 0.2235 (n.s) |
| | T-test | WT, PDS_GFP | T=-0.60715, df=14 | 0.5535 (n.s) |
| Stigmasterol | Lavene's test | WT, PDS_GFP | F=1.6604 | 0.2184 (n.s) |
| | Shapiro-Wilks test | WT, PDS_GFP | W=0.95485 | 0.5702 (n.s) |
| | T-test | WT, PDS_GFP | T=1.6733, df=14 | 0.1165 (n.s) |
| isofucosterol | Lavene's test | WT, PDS_GFP | F=2.6682 | 0.1246 (n.s) |
| | Shapiro-Wilks test | WT, PDS_GFP | W=0.96981 | 0.8356 (n.s) |
| | T-test | WT, PDS_GFP | T=-1.9239, df=14 | 0.07494 (n.s) |
| B-Sitosterol | Lavene's test | WT, PDS_GFP | F=13.177 | 0.002731 (**) |
| | Shapiro-Wilks test | WT, PDS_GFP | W=0.95923 | 0.6479 (n.s) |
| | Mann-Whitney U test | WT, PDS_GFP | W=32 | 1.00 (n.s) |

**Supplementary Table 6. Table of Statistics for Supplementary Figure 4.**
*=P<0.05, ** P<0.01,*** P<0.0001

| Experiment | Test | Sample | Value | P-Value |
|---|---|---|---|---|
| *PDS1* expression in leaf for WT, PDS_GFP and PDS_CAS | Lavene's test | WT, PDS_GFP, PDS_GFP_FT | F=1.8661 | 0.1807 (n.s) |
| | Shapiro-Wilks test | WT, PDS_GFP, PDS_GFP_FT | W=0.85933 | 0.004035 (**) |
| | Kruskal-Wallis | WT, PDS_GFP, PDS_GFP_FT | $X^2$=1.6238 | 0.444 (n.s) |
| *PDS2* expression in leaf for WT, PDS_GFP and PDS_CAS | Lavene's test | WT, PDS_GFP, PDS_GFP_FT | F=1.4786 | 0.2518 |
| | Shapiro-Wilks test | WT, PDS_GFP, PDS_GFP_FT | W=0.92823 | 0.1002 |
| | One-way ANOVA | WT, PDS_GFP, PDS_GFP_FT | F=4.323 | 0.0275 (*) |
| | Post-hoc Tukey HSD | WT-PDS_GFP | 0.0242939 (*) | |
| | | WT-PDS_GFP:FT | 0.5779040 (n.s) | |
| | | PDS_GFP-PDS_GFP:FT | 0.1567101 (n.s) | |

**Supplementary Table 7. List of *Agrobacterium* Strains.**

| **Strains** | **Chromosomal background** | **Ti plasmids** | **Resistance gene** |
|---|---|---|---|
| AGL1 | C58, *RecA* | pEHA105 (pTiBo542DT-DNA) | rif, carb |
| LBA4404 | *Ach5* | pAL4404 | rif, strep |
| GV3101 | C58 | pMP90 (pTiC58DT-DNA) | rif, gent |

**Supplementary Table 8. Table of Primers.**

| **Cloning Primers** | | | |
|---|---|---|---|
| **Target gene** | **Forward primer (5'-3')** | **Reverse Primer (5'-3')** | **Amplicon Length** |
| GCTT_*TXSS*_GGTG | TGTGGTCTCTGCTTCTCTTGTAACTCAAGC | ACAGGTCTCTACTAGGACATTCACAGCATC | 328 |
| GCTT_*TXSS*_TAGT | TGTGGTCTCTGCTTCTCTTGTAACTCAAGC | ACAGGTCTCTCACCGGACATTCACAGCATC | 328 |
| GCTT_*CAS*_ GGTG | TGTGGTCTCTGCTTCCTGAGATGTGGC | ACAGGTCTCTCACCATGGGCTCCAC | 312 |
| TATG_*PDS*_GCTT | TGTGGTCTCTTATGGAAGCAAGAGACG | ACAGGTCTCTAAGCTTTCTCAGGCC | 319 |

| | | | |
|---|---|---|---|
| TAGT_*FT*_ GGTG | TGTGGTCTCTTAGTCTATAAATATAAGAGAtCC | ACAGGTCTCTCACCTTGGCCATAAGTAACC | 126 |
| GCTT_*GFP*_ GGTG | TGTGGTCTCTGCTTTGACCACCTTCAGCTACGG | ACAGGTCTCTCACCTGCCGTTCTTCTGCTTGTCG | 328 |
| GCTT_*GFP*_ GGTG | TGTGGTCTCTGCTTTGACCACCTTCAGCTACGG | ACAGGTCTCTACTATGCCGTTCTTCTGCTTGTCG | 328 |
| MVP | ATGGAAGACAAGTCATTGG | TTAAGACGAGTTTTTCTTATTAGG | 759 |
| **qPCR Primers** | | | |
| **Target gene** | **Forward primer (5'-3')** | **Reverse Primer (5'-3')** | **Amplicon Length** |
| PP2A.2 | CATCGGTGTATGGTCCCGTTA | CTTTCTGCCACCTGAAAATGCA | 74 |
| SAND.2 | TCTTTCAGTTGGAACCCTGCA | CTGCAATATAGCACCAGCAGC | 93 |
| CoPDS1.1 | GGGAAGTGGAAGTGGTTCCT | AGTGGCACTGCTATGAGGTT | 74 |
| CoPDS2.1 | ATTTCAATTCCGTTCTTCTG | TCGGAAGGTTGAAGACAATA | 139 |
| CoCAS2.1 | GGAGACTTCCCACAACAGGA | AACAAGGTGGCTGAAGGACT | 133 |
| CoCAS4.2 | GGCAATGGAAAAGGGACGAC | GCCACATTTCAGGGGGTAAT | 136 |

**Supplementary Table 9. Table of Statistics for Outliers.**
*=P<0.05, ** P<0.01,*** P<0.0001

| Experiment | Test | Sample | Value | P-Value |
|---|---|---|---|---|
| Stigmasterol metabolite accumulation in leaf | Grubbs test | PDS_CAS.2 | G=2.13833 U=0.25347 | 0.02265 (*) |
| *PDS1* expression in leaf for WT, PDS_GFP and PDS_CAS | Grubbs test | WT.4 | G=2.32214 U=0.11962 | 0.002243 (**) |
| *PDS2* expression in leaf for WT, PDS_GFP and PDS_CAS | Grubbs test | WT.4 | G=2.27371 U=0.15595 | 0.005048 (**) |
| *CAS2* expression in leaf for WT, PDS_GFP and PDS_CAS | Grubbs test | WT.4 | G=2.26720 U=0.16078 | 0.005543 (**) |
| *PDS1* expression in Flower for WT, PDS_GFP and PDS_CAS | Grubbs test | WT.2 | G=2.03721 U=0.32241 | 0.04823 (*) |

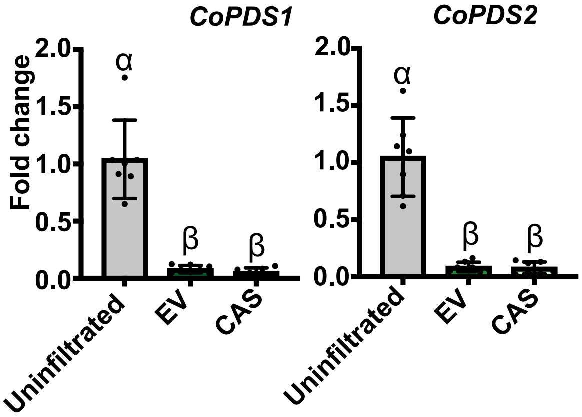


**Supplementary Figure 1. Virus induced gene silencing of phytoene desaturase in comparison to uninfiltrated plant in leaves.** Expression levels of CoPDS genes in pot marigold leaves 38 days post infiltraton with VIGS vectors as determined by qRT-PCR; n ≥ 7. Statistical significance was determined by a Kruskal-Wallis test with a post-hoc Dunn test into groupings represented with α or β (Supplemental Table 3)

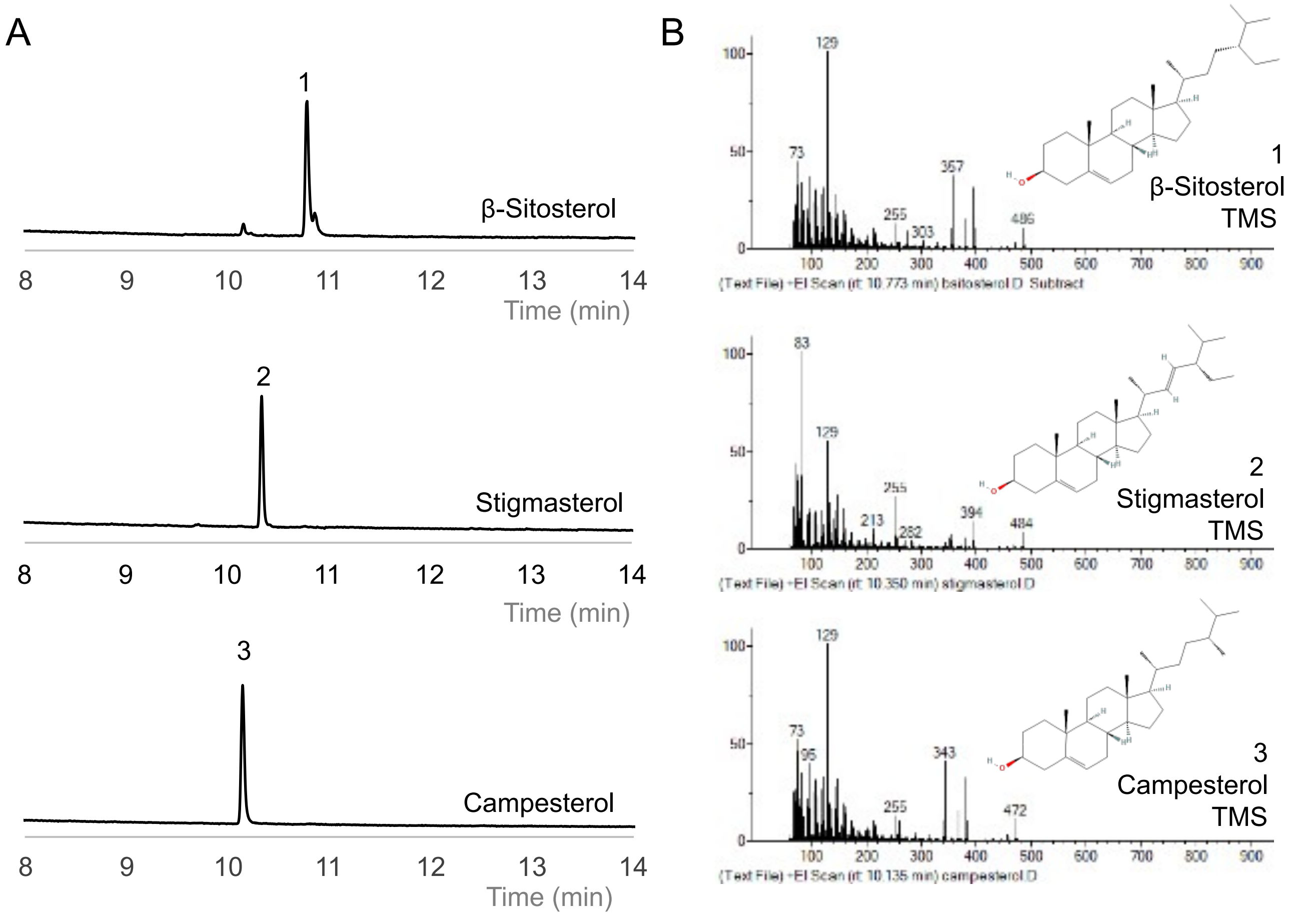


**Supplementary Figure 2. GC-MS of sterol standards and associated mass spectra.** (A) GCMS Total Ion chromatogram (TIC) of trimethylsilyl (TMS) derivatised sterols including 1. B-sitosterol, 2. stigmasterol, 3. campesterol. Sterols were kindly provided by Anne Osbourn, John Innes Center, Norwich, UK. (B) Mass spectra for TMS derivatised sterols

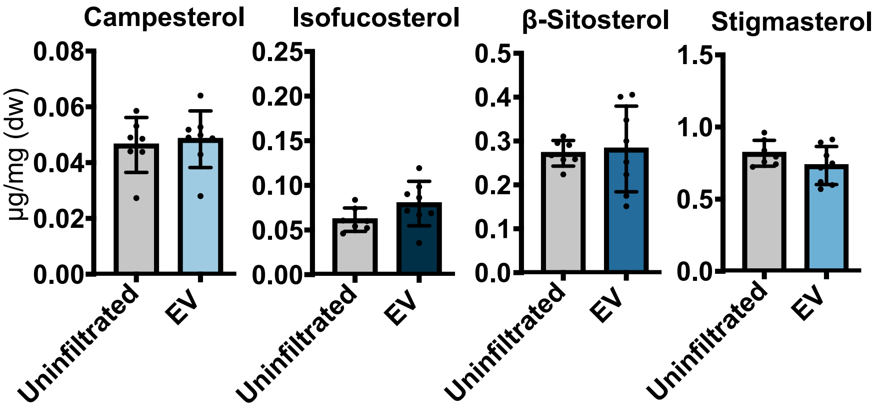


**Supplementary Figure 3. Sterol accumulation did not change after virus induced gene silencing of *phytoene desaturase* in comparison to uninfiltrated plant in leaves.** Quantification of campesterol, isofucosterol, β-sitosterol and stigmasterol in plants with reduced *CoPDS* expression compared to uninfiltrated plants as determined by GCMS compared to an internal standard; n ≥ 7. statistical significance was calculated using a students t-test for campesterol, isofucosterol and stigmasterol. While a Mann-Whitney U test was used to test for significance in β-sitosterol as it was not of equal variance (supplemental Table 5); EV = empty vector (pTRV2-PDS:GFP).

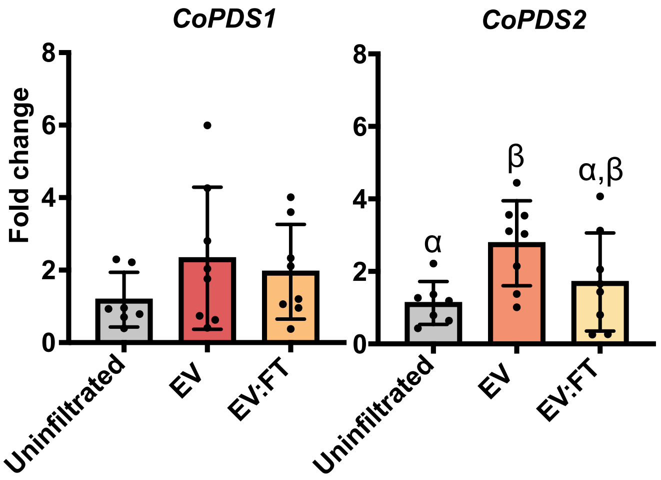


**Supplementary Figure 4. Virus induced gene silencing of *phytoene desaturase* in comparison to uninfiltrated plant in flowers.** Expression levels of *CoPDS* genes in pot marigold flowers 66 days post infiltraton with VIGS vectors as determined by qRT-PCR; n ≥ 7. Statistical significance was determined by a Kruskal-Wallis test for *CoPDS1* as the data was not normally distributed and ANOVA with post-hoc Tukey test for *CoPDS2* into groupings represented with α or β (Supplemental Table 6); EV = empty vector (pTRV2-PDS:GFP); EV:FT = empty vector *flowering locus T* fusion (pTRV2-PDS:GFP:FT).
